# Characterisation of key genotypic and phenotypic traits of clinical cystic fibrosis *Staphylococcus aureus* isolates

**DOI:** 10.1101/2022.12.20.520977

**Authors:** Micaela Mossop, Luca Robinson, Jhih-Hang Jiang, Anton Y. Peleg, Luke V. Blakeway, Nenad Macesic, Audrey Perry, Stephen Bourke, Fatima R. Ulhuq, Tracy Palmer

## Abstract

**Introduction:** One third of people with CF in the UK are co-infected by both *Staphylococcus aureus* and *Pseudomonas aeruginosa*. Chronic bacterial infection in CF contributes to the gradual destruction of lung tissue, and eventually respiratory failure in this group.

**Gap Statement:** The contribution of *S. aureus* to cystic fibrosis (CF) lung decline in the presence or absence of *P. aeruginosa* is unclear. Defining the molecular and phenotypic characteristics of a range of *S. aureus* clinical isolates will help further understand its pathogenic capabilities.

**Aim:** Our objective was to use molecular and phenotypic tools to characterise twenty-five clinical *S. aureus* isolates collected from mono- and coinfection with *P. aeruginosa* from people with CF at the Royal Victoria Infirmary, Newcastle upon Tyne.

**Methodology:** Genomic DNA was extracted and sequenced. Multilocus sequence typing was used to construct phylogeny from the seven housekeeping genes. A pangenome was calculated using Roary. and cluster of Orthologous groups were assigned using eggNOG-mapper which were used to determine differences within core, accessory, and unique genomes. Characterisation of sequence type, clonal complex, *agr* and *spa* types was carried out using PubMLST, eBURST, AgrVATE and spaTyper, respectively. Antibiotic resistance was determined using Kirby Bauer disk diffusion tests. Phenotypic testing of haemolysis was carried out using ovine red blood cell agar plates and mucoid phenotypes visualised using Congo red agar.

**Results:** Clinical strains clustered closely based on *agr* type, sequence type and clonal complex. COG analysis revealed statistically significant enrichment of COG families between core, accessory and unique pangenome groups. The unique genome was significantly enriched for replication, recombination and repair, and defence mechanisms. The presence of known virulence genes and toxins were high within this group, and unique genes were identified in 11 strains. Strains which were isolated from the same patient all surpassed average nucleotide identity thresholds, however, differed in phenotypic traits. Antimicrobial resistance to macrolides was significantly higher in the coinfection group.

**Conclusion:** There is huge variation in genetic and phenotypic capabilities of *S. aureus* strains. Further studies on how these may differ in relation to other species in the CF lung may give insight into inter-species interactions.

**Data summary:** The assembled GenBank (gbk) files for all clinical isolates in this study have been deposited in ENA under the study accession PRJEB56184, accession numbers for each of the twenty-five clinical isolates have been provided in Table S1. The reference strains were collected from the NCBI BioSample database (www.ncbi.nlm.nih.gov/biosample): MRSA_252 (NC_002952.2), HO 5096 0412 (NC_017763.1), ST398 (NC_017333.1) and NCTC8325 (NC_007795.1).

## INTRODUCTION

In the UK 1 in every 2,500 babies is born with a mutation in both copies of the *cystic fibrosis transmembrane regulator* (*CFTR*) gene. Pathogenic mutations cause a disruption in epithelial bicarbonate and chloride secretion and hyperabsorption of sodium, leading to an osmotic imbalance that results in thick and sticky mucus build-up in the lungs, which is characteristic of cystic fibrosis (CF) (1). Within the airways, the nutritionally rich environment traps bacteria and promotes their growth, while clearance via the ciliary escalator is impaired. Infection contributes to the continuous inflammation of the airways and leads to the gradual destruction of lung tissue. As lung infection shapes the clinical outcome of individuals and is the leading cause of mortality in people with CF (2), it is an area of intense medical research.

Two of the most prevalent bacterial species in CF pulmonary infections are *Staphylococcus aureus* and *Pseudomonas aeruginosa*. Paediatric CF patients are often colonised with *S. aureus* from an early age (<1 year old). To prevent this, in the UK, CF patients are prescribed prophylactic flucloxacillin from birth or point of diagnosis (2). As patients reach adulthood, *P. aeruginosa* typically overtakes *S. aureus* and becomes the dominant species in the lungs. However, both species are co-isolated from one third of UK adults with CF (2). Some studies suggest that coinfection with both species correlates with severe lung function decline and worsened patient clinical outcomes compared to patients that are monoinfected by either pathogen (3–5). However, it has also been found that *P. aeruginosa* infection results in poor patient outcome independent of the presence of *S. aureus* (6). Most evidence suggests *S. aureus* infections in CF tend to be a marker of milder disease compared to patients infected with *P. aeruginosa*, particularly in adults, with these patients exhibiting higher lung function, and fewer exacerbation periods (6, 7). By contrast, other work has shown that *S. aureus* has persistence capabilities similar to *P. aeruginosa* and patients with high *S. aureus* density in throat cultures exhibit rapid lung function decline (8). To date, no consensus has been reached in the literature regarding the significance of *S. aureus* in the presence or absence of *P. aeruginosa*, and recent reviews highlight the need to investigate clinical isolates further (9, 10).

Much of the previous work has helped to define the mechanisms that *P. aeruginosa* employs to chronically infect the CF lung, including loss of motility, a lack of a functional type III secretion system, and conversion to mucoid colony phenotype (11). A growing number of studies have also started to elucidate how *S. aureus* adapts to long-term colonisation of the CF lung. These include mutations in metabolic genes, resulting in persistent small colony variants (SCVs) and in regulatory genes, such as accessory gene regulator (*agr*) mutants, which present an inactive quorum-sensing *agr* system, and therefore reduced expression of virulence factors including haemolysins (12–14). Increased extracellular polysaccharide (EPS) production facilitates the biofilm mode of growth that also contributes to persistence and antibiotic recalcitrance in the lungs (15). Furthermore, bacteriophages become mobilised with a higher frequency in infections, raising the incidence of mobile genetic elements, which include known toxins and antimicrobial resistance (AMR) genes (16).

It is generally accepted that CF lung infection is polymicrobial, and an understanding of lung microbiome composition and interspecies interactions is critical to understand disease progression. Although historically CF pathogens have been studied in isolation, research has shown that competitive and cooperative interactions between *S. aureus* and *P. aeruginosa* can alter survival and persistence in the CF lung for example through increased antimicrobial resistance profiles and improved immune system evasion (16). Further work is required to understand the molecular basis for cooperative and competitive interactions in the CF lung and how these influence pathogeneses.

In this study we collected and characterised 25 clinical *S. aureus* isolates from 21 chronically infected adult CF patients from the Royal Victoria Infirmary (RVI, Newcastle-upon-Tyne), including determining fully assembled genome sequences. The *S. aureus* isolates belonged to two groups, (i) from patients that had never cultured *P. aeruginosa* since their first visit to the clinic (monoinfected patients, n=10) and (ii) from patients that had also previously isolated *P. aeruginosa* (coinfected patients, n=11). Isolates were selected from these two categories with the aim of identifying whether there were genomic signatures associated with *S. aureus* isolates from the two different infection groups. Interestingly, we noted different *S. aureus* phenotypic behaviour from isolates within the same patient, despite genome analysis indicating they are the same strain, showcasing the heterogeneity of *S. aureus* phenotypes within the same CF lung. A significantly higher proportion of coinfection isolates were resistant to macrolides in CF patients, particularly those chronically infected by *P. aeruginosa*.

## METHODS

### Clinical isolates and patient data

Respiratory samples were collected from patients attending the Adult Cystic Fibrosis Centre as part of their standard clinical care. Patients gave signed informed consent for the collection of clinical data for research and inclusion in the national CF registry, with approval from the regional ethics committee. Patients were classed as monoinfected if they had only cultured *S. aureus* from the time of first visit to the clinic. Coinfected patients had produced positive cultures for both *S. aureus* and *P. aeruginosa* from the time their records began.

For lung function assessments, patients performed spirometry as part of their clinical outpatient visit according to the ERS/ATS guidelines (17). Percent predicted values were generated for the Forced Expiratory Volume of air in the first second (FEV_1_) using the Global Lung Initiative reference equations (18). All other demographic information was obtained from hospital records.

### DNA extractions

Clinical isolates were cultured on Tryptone soya agar (TSA, Oxoid) at 37°C. Genomic DNA (gDNA) was extracted using the GenElute Bacterial Genomic DNA Kit (Sigma Aldrich) according to the manufacturer’s instructions.

### Genome assembly and annotation

Short read NGS libraries were prepared using the Nextera flex DNA Library Prep Kit (Illumina), and 150 bp paired-end sequencing was performed on the NovaSeq 6000 system. Read trimming and quality control was performed using Trim Galore (v0.6.5) (http://www.bioinformatics.babraham.ac.uk/projects/trimgalore/) and assembly was performed using Unicycler (v0.4.8) (19), with default parameters. Alignments were performed against the reference genome *S. aureus* NCTC 8325 (NC_007795.1) (20) using progressive Mauve (21), with default parameters. The 25 genomes were annotated using Prokka (Prokaryotic genome annotation) (v1.14.5) with a similarity cut off e-value 1e-6 and minimum contig size of 200 bp (22).

### Phylogenetic analysis

Nucleotide sequences for the seven housekeeping genes (*arcC, aroE, glpF, gmk, pta, tpi*, and *ygiL*) were extracted from the assembled genomes of the 25 clinical *S. aureus* isolates and four laboratory/reference strains of *S. aureus:* MRSA_252 (NC_002952.2), HO 5096 0412 (NC_017763.1), ST398 (NC_017333.1) and NCTC8325 (NC_007795.1). Nucleotide sequences were concatenated and aligned using MUSCLE (default parameters) (23, 24) and a maximum likelihood tree for nucleotide sequences was constructed in MEGA X (25) with Hasegawa-Kishino-Yano model, Gamma distribution with invariant sites G+I, and bootstrapping (n = 1000) parameters. The tree was midpoint rooted and visualised using the iTOL webserver (26).

### Pangenomic analysis

Based on the GFF3 files produced by Prokka (22), a pangenome was calculated using Roary (v.13.3.0) (27) with default parameters and a minimum percentage identity of 75% between predicted protein homologues. Functional sequence annotation by orthologs to establish COG categories for the core, accessory, and unique genes was performed using eggNOG-mapper v5 (28), with default parameters. The percentage conservation of a gene within the pangenome to be classed as (i) core was 95%, ≥ 28 strains, (ii) accessory, 5-95%, 2-27 strains, or (iii) unique, only one strain. Statistical analysis was conducted using X^2^ test with a critical value of *p* = 0.00128205 after Bonferroni correction (39 tests were conducted = 13 COG groups x 3 pangenome groups). Statistical analysis was only carried out for COG categories where at least 5 genes per pangenome group were annotated by eggNOG mapper.

### Sequence analysis

Sequence type was determined using PubMLST (29) [last accessed 12.08.2022]. Sequences were grouped into clonal complexes using the eBURST algorithm (30). The *agr* type was assigned using AgrVATE (v1.0.2) (31) and *spa* type using spaTyper (v0.3.1) (32). Average nucleotide identities (ANI) were calculated using fastANI (33). Default parameters were used throughout.

Antimicrobial resistance predictions were obtained using both MYKROBE (v0.9.0) (34) as well as ResFinder v4.1 (35), with default parameters. To determine the presence/absence of toxins/virulence genes, assembled genomes were aligned using Artemis Comparison Tool (ACT) (36). Multiple sequence alignment of the protein sequences was performed using ClustalW (v2.1) (37) and visualised using Jalview v2 (38), with default parameters.

### Statistical analysis

All statistical tests were carried out using GraphPad Prism (v 9.0) except for X^2^ tests which were calculated on Microsoft Excel.

### Bacterial strains and growth conditions

*S. aureus* isolates were grown at 37°C in Tryptone soya broth (TSB, Oxoid) and plated on Tryptone soya agar (TSA, Oxoid). Antimicrobial susceptibility testing was performed on Mueller-Hinton agar (MHA, Oxoid) plates.

### Haemolysis assay

*S. aureus* haemolysis was assayed by spotting 2 μl of *S. aureus* cultures grown in TSB (37 °C, 15 hrs) on commercial sheep blood agar plates. Plates were incubated for 15 hrs at 37 °C followed by a 4°C incubation step for a further 24 hrs to detect hot-cold activation of β-haemolysin before imaging (39). *S. aureus* COL and RN6390 were used as negative and positive controls, respectively.

### Polysaccharide production assay

Sterile Congo red dye stock solution contained 5 g of Congo red (Sigma Aldrich) in 100 ml of distilled water. A combination of 11.1 g Brain Heart Infusion Broth (Sigma Aldrich), 15 g sucrose (Fisher Scientific) and 3 g select agar (Life Technologies) were added to 300 ml of distilled water and autoclaved. After cooling to ~ 50 °C, 4.8 ml of 5% Congo red stock solution was added to make the Congo red agar (CRA) plates. Single *S. aureus* colonies were picked from TSA plates and grown in TSB for 15 hrs at 37 °C. Then, 2 μl of overnight stationary cultures were spotted on freshly prepared CRA plates. CRA plates were incubated for 15 hrs at 37°C and imaged.

### Antimicrobial susceptibility testing

Antimicrobial susceptibility testing was performed using the Kirby Bauer disk diffusion test. Several *S. aureus* colonies were picked, transferred to 5 ml TSB and mixed until a homogenous suspension was reached. The OD_600_ was adjusted to OD_600_ 0.125 (equivalent to McFarland Standard 0.5). A sterile cotton swab was used to distribute the culture evenly over the surface of a Mueller-Hinton agar plate and left to dry at room temperature for 5-10 mins. Antibiotic disks (no more than six per plate) were applied in the appropriate formation: cefoxitin (FOX) (30 μg), chloramphenicol (C) (30 μg), ciprofloxacin (CIP) (5 μg), clindamycin (DA) (2 μg), erythromycin (E) (15 μg), rifampicin (RP) (5 μg), penicillin (PG) (1 μg) and tetracycline (TE) (30 μg), were dispensed. The inoculated plate was then incubated at 35 °C, for 18-24 hours and zones of inhibition were measured across the full diameter. Breakpoints taken from EUCAST v12.0.

## RESULTS

### Clinical isolates and patient data

To investigate the genetic diversity of the *S. aureus* strains, 25 *S. aureus* isolates (SA-1 – SA-21) were collected and sequenced from 21 CF patients as outlined in methods. In total 11 *S. aureus* strains were collected from ten adult CF patients that were colonised by *S. aureus* only. Additionally, for comparison, 14 *S. aureus* isolates from 11 adult CF patients who had previously cultured both *S. aureus* and *P. aeruginosa* were also characterised. Four patients (one monoinfected and three co-infected) provided two separate isolates termed a and b **(Table 1)**. For isolates numbered six and ten these a and b samples were collected on two separate visits to the clinic whereas for isolates numbered three and seven they were collected at the same visit but represented two different colony types. SA-11 was classified as a thymidine dependent small-colony variant (SCV) upon streaking and growth was only achieved in on Mueller-Hinton agar in the presence of 5% CO_2_.

**Table 1.** Strain and patient metadata. *S. aureus* strains are labelled numerically SA-1 – SA-21. Samples labelled A and B were isolated from the same patient. Orange labels indicates these strains are from the *S. aureus* monoinfection group and blue labels indicate strains were isolated from coinfections with *P. aeruginosa*. Patient sex is classified as M (male) or F (female). FEV_1_ results are given as exact value in litres followed by FEV_1_ percent predicted. Body mass index given as kg/m^2^. Abbreviations for oral and inhaled antibiotics are Azithromycin (Az), Flucloxacillin (Flu), Colistin (Col) and Tobramycin (Tob). Co-morbidity abbreviations are pancreas sufficient (PS) pancreas insufficient (PI), CFRD (CF-related diabetes), renal calculi (RC), gastroesophageal reflux disease (GORD), intestinal obstruction (DIOS), aspergillosis (ABPA), nasal polyps (NP). Boxes populated with (-) indicate the patient was not on any antibiotic regime and/or presented no co-morbidities. Boxes populated with (“) indicate the information is the same as the row directly above.

| Strain ID | Sample date | Sample type | Age | Sex | FEV<sub>1</sub> | BMI | CFTR mutation | Oral Abx | Inh Abx | Co-morbidities |
| --- | --- | --- | --- | --- | --- | --- | --- | --- | --- | --- |
| ***S. aureus* mono-infection strains** |  |  |  |  |  |  |  |  |  |  |
| SA-1 | 03/07/19 | Cough swab | 37 | M | 3.18<br>79% | 25.6 | F508del/<br>R334W | - | - | PS |
| SA-3A | 12/07/19 | Sputum | 68 | F | 1.78<br>78% | 33.7 | F508del/<br>P67L | Az | - | PI, CFRD |
| SA-3B | 12/07/19 | " | " | " | " | " | " | " | " | " |
| SA-8 | 14/08/19 | Sputum | 32 | F | 2.82<br>85% | 21.1 | F508del/<br>R117H-5T | Flu | - | PS |
| SA-11 | 12/08/19 | Sputum | 22 | F | 3.26<br>87% | 22.9 | F508del/<br>P67L | Az | - | PS |
| SA-12 | 23/08/19 | Cough swab | 24 | F | 3.31<br>93% | 22.6 | F508del/<br>G551D | Flu | - | PS, CFRD, RC |
| SA-13 | 23/08/19 | Sputum | 34 | F | 2.34<br>69% | 24.6 | F508delA<br>rg1066His | Flu | - | PS, RC |
| SA-14 | 20/09/19 | Cough swab | 21 | F | 2.38<br>69% | 20.1 | F508del/<br>F508del | Flu | Col | PI |
| SA-16 | 08/11/19 | Cough swab | 25 | M | 4.22<br>85% | 30.5 | F508del/<br>R117H-5T | - | - | PS |
| SA-17 | 15/11/19 | Cough swab | 65 | M | 2.04<br>68% | 29.5 | F508del/<br>Q1291H | - | - | PI, CFRD, NP |
| SA-21 | 03/01/20 | Cough swab | 28 | M | 3.18<br>83% | 24.1 | F508del/<br>F508del | Flu | - | PI, DIOS |
| ***S. aureus* co-infection strains** |  |  |  |  |  |  |  |  |  |  |
| SA-2 | 12/07/19 | Cough swab | 24 | F | 1.86<br>59% | 23.5 | G542X/<br>A455E | Az<br>Flu | - | PI |
| SA-4 | 12/07/19 | Sputum | 36 | F | 3.2<br>110<br>% | 25 | F508del/<br>I507del | Az | - | PI, CFRD, GORD |
| SA-5 | 17/07/19 | Sputum | 50 | F | 1.42<br>56% | 25.1 | F508del/<br>R117H-5T | Flu | Col | PS |
| SA-6A | 29/07/19 | Sputum | 28 | M | 1.45<br>34% | 19.1 | F508del/<br>F508del | Az<br>Flu | Col | PI |
| SA-6B | 14/11/19 | " | " | " | " | " | " | " | " | " |
| SA-7A | 19/07/19 | Sputum | 36 | M | 2.06<br>50% | 19 | F508del/<br>F508del | Az<br>Flu | Col | PI, CFRD, ABPA |
| SA-7B | 19/07/19 | " | " | " | " | " | " | " | " | " |
| SA-9 | 12/08/19 | Sputum | 21 | F | 2.3<br>76% | 22.1 | F508del/<br>G85E | Flu | - | - |
| SA-10A | 14/08/19 | Cough swab | 27 | M | 2.05<br>73% | 25.6 | P67L/<br>G551D | Flu | - | PS, ABPA |
| SA-10B | 14/11/19 | " | " | " | " | " | " | " | " | " |
| SA-15 | 04/10/19 | Cough swab | 36 | M | 3.1<br>73% | 23.8 | G551D/T<br>rp401Ter | - | - | PI, NP |
| SA-18 | 11/12/19 | Cough swab | 39 | M | 0.89<br>26% | 23.8 | F508del/<br>F508del | Flu | Col | PI, CFRD, NP |
| SA-19 | 11/12/19 | Sputum | 20 | F | 2.68<br>86% | 28.6 | G551D/<br>Y1032C | - | - | PI |
| SA-20 | 08/01/20 | Cough swab | 19 | F | 3.05<br>95% | 28.6 | F508del/<br>R347H | Flu | Col/<br>Tob | PS |

The isolates were obtained from nine male and 12 female patients with a median age of 28 (range, 19 to 68 years), and a variety of CFTR mutations **(Table 1)**. When comparing health indicators between the two groups, the mean FEV_1_% of patients with *S. aureus* monoinfections was 12.5% higher than coinfected patients (79.6% vs 67.1%), indicating better lung function, however, this was not statistically significant using a t-test (*p* = 0.15). No significant difference was noted between the mean BMI (body mass index kg/m^2^) values (25.47 monoinfected vs 24.0 coinfection) and no observable co-morbidity patterns were found between the two groups.

A phylogenetic tree of all 25 clinical *S. aureus* isolates, and four non-CF associated laboratory strains (NCTC8325, ST398, HO_506_0492, and MRSA252) was constructed to determine the genetic relationship between strains **(Figure 1)**. Further analysis showed that the strains collectively represent 13 sequence types (ST), the most common was ST 5 with six clinical isolates belonging to this group **(Figure 1)**. STs were then assigned into seven clonal complexes (CC) using eBURST (40). Strains of the same CC and *agr* type clustered together, and in general, strains also clustered on the same branch depending on whether they were isolated from mono- or coinfections. However, there were some exceptions, SA-13, which is a monoinfection strain that shares a branch alongside coinfection strains and SA-6A and SA-6B which were isolated from a co-infected patient and cluster near strains isolated from patients who were *P. aeruginosa* culture negative.

**Figure 1.**
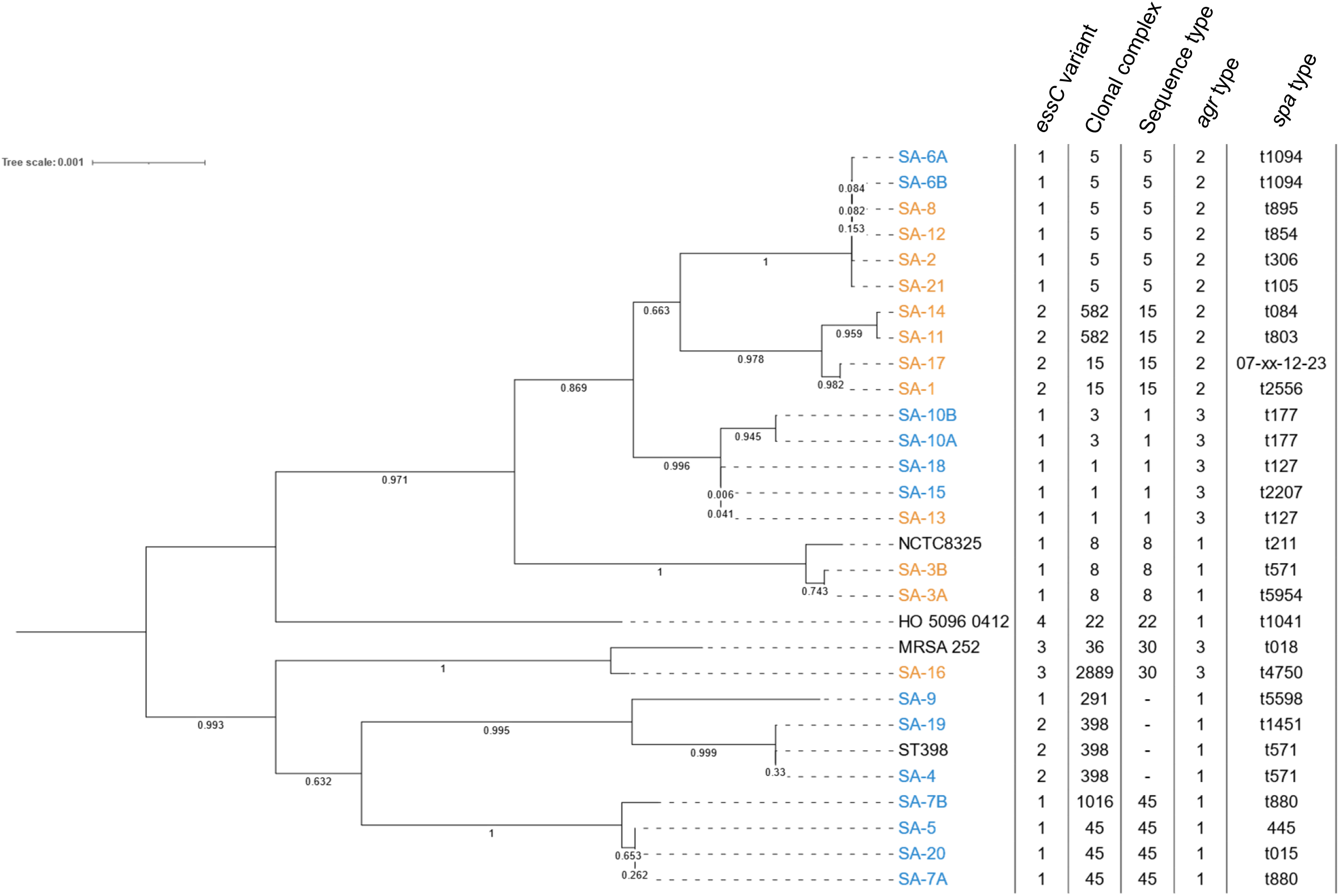
Phylogenetic tree of clinical *S. aureus* isolates. Phylogenetic tree constructed based on the 7 housekeeping genes (*arcC, aroE, glpF, gmk, pta, tpi, and yqiL*) from the 25 clinical *S. aureus* isolates and 4 laboratory strains of *S. aureus* using the maximum likelihood method in MEGAX with Hasegawa-Kishino-Yano model, Gamma distribution with invariant sites G+I, and bootstrapping (n = 1000) parameters. Strains isolated from mono-infections are labelled in orange, strains from co-infections are labelled in blue and reference strains are labelled in black. Sequence type was determined using PubMLST, clonal complexes were determined using eBURST. The *agr* type was assigned using AgrVATE and *spa* type using spaTyper. Tree editing was carried out using iTOL.

The *S. aureus* Type VII secretion system (T7SS) has been shown to play a role in interbacterial competition through the secretion of protein toxins (41, 42). The T7SS occurs as one of four variants, depending on the sequence of the *essC* gene, which encodes a core component of the secretion system (43, 44). One of the key differences between all four variants is some of the toxins secreted by the T7SS, with at least one variant-specific toxin found for each of the four *essC* subtypes (41, 43, 44). Most of the clinical isolates had the most common variant, *essC1*, with *essC2* found in seven isolates and *essC3* in only one (Figure 1). T7SS *essC4* appears to be largely restricted to CC22 (44) and no *essC4* variants were found among the clinical strains. While the T7SS is not essential for *S. aureus* viability, it has been reported to be conditionally essential for survival in the presence of *P. aeruginosa* in a chronic wound infection model (45). As shown in Fig 1, all but two of the coinfection isolates were *essC1* variants, whereas the mono-infection strains had a more even split between *essC1* and *essC2* subtypes. However, at present the relevance of this, if any, is unclear since none of the T7 secreted toxins identified to date have been shown to target *P. aeruginosa*.

### *S. aureus* genomic characterisation

The pangenome is the entire collection of genes present within a species and can be divided into three groups (i) the core genome, which is shared by all strains (95%, ≥ 28 strains), (ii) the accessory genome, consisting of genes present in some but not all genomes (5-95%, 2-27 strains) and (iii) the unique genome, composed of genes (singletons) present in only one strain (3.4%). We estimated the pangenome to be made up of 4,315 orthologs, with a split of (i) core as 2,070 (47.9 %), (ii) accessory as 1,564 (36.2 %), and (iii) 681 singletons (15.8 %) **(Figure 2)**. From these genes, a total of 3,534 (81.9%) were identified and assigned cluster of orthologous groups (COG) category using the eggNOG-mapper (28). **Figure 2** shows the distribution of the COG categories within the core, accessory, and unique genomes.

**Figure 2.**
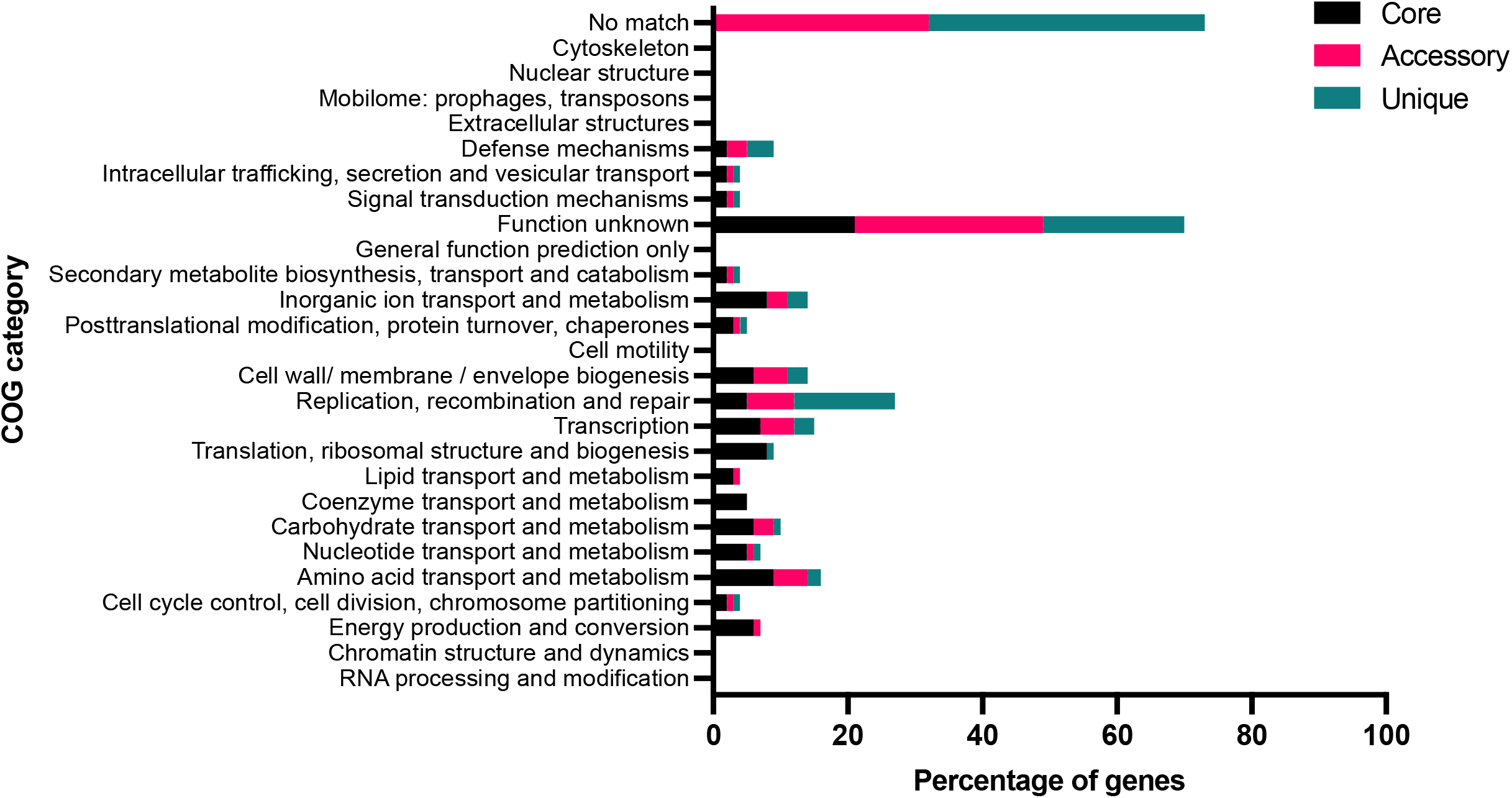
COG classifications for core, accessory, and unique genes present in *S. aureus* clinical isolates. Bar chart represents the percentage of genes in the core, accessory and unique genes which were assigned a known COG function by the eggNOG-mapper v5 software.

After assignment of COG groups to the genes belonging to the core, accessory and unique genomes, the enrichment of COG families was investigated within the pangenome groups. As expected, the core genome was enriched for all COG families related to metabolism, as well as translation, ribosomal structure and biogenesis, and transcription, many of which are essential genes **(Table 2)**. Interestingly, the unique genome was significantly enriched for the replication, recombination and repair, and defence mechanism COG families **(Table 2)**.

**Table 2.** Enrichment of COG categories within the core, accessory, and unique genomes. COG categories were assigned using eggNOG-mapper v5. Statistical tests were carried out using X^2^ on categories where at least 5 genes per pangenome group had been annotated. This statistical test assesses the likelihood of significant enrichment in the different COG categories between the core, accessory, and unique groups. The *p*-value given has been adjusted using the Bonferroni correction, (*) highlights significant *p*-values.

| COG Letter | COG category | Core | Accessory | Unique | *p*-value |
| --- | --- | --- | --- | --- | --- |
| Percentage of genes with COG annotation |  |  |  |  |  |
| **D** | Cell cycle control, cell division, chromosome partitioning | 1.79% | 0.90% | 1.32% | 0.074 |
| **E** | Amino acid transport and metabolism | 9.23% | 5.18% | 2.20% | <0.001* |
| **F** | Nucleotide transport and metabolism | 4.54% | 0.77% | 0.73% | <0.001* |
| **G** | Carbohydrate transport and metabolism | 6.43% | 2.81% | 1.32% | <0.001* |
| **J** | Translation, ribosomal structure, and biogenesis | 8.02% | 0.34% | 0.29% | <0.001* |
| **K** | Transcription | 7.20% | 5.05% | 3.08% | <0.001* |
| **L** | Replication, recombination, and repair | 4.64% | 6.78% | 14.83% | <0.001* |
| **M** | Cell wall/ membrane / envelope biogenesis | 5.70% | 5.12% | 2.79% | 0.0104 |
| **P** | Inorganic ion transport and metabolism | 7.78% | 2.94% | 2.50% | <0.001* |
| **S** | Function unknown | 20.97% | 28.39% | 21.00% | <0.001* |
| **T** | Signal transduction mechanisms | 1.74% | 1.47% | 0.88% | 0.277 |
| **U** | Intracellular trafficking, secretion, and vesicular transport | 1.69% | 0.90% | 0.88% | 0.065 |
| **V** | Defence mechanisms | 1.50% | 3.20% | 4.41% | <0.001* |

The unique genes present in each clinical *S. aureus* isolate were then further examined. Many were found to be unique mobile genetic elements (MGEs). The genetic plasticity of the *S. aureus* genome is largely attributed to MGEs, which include prophages, plasmids, pathogenicity islands (SaPIs), staphylococcal cassette chromosome *mec* (SCCmec), and transposons. MGEs can disseminate between bacteria through horizontal gene transfer (HGT) mechanisms and are particularly concerning as they can alter virulence and resistance profiles (46, 47).

Evidence of phage integration was present across the genomes of all *S. aureus* clinical strains, although the phage-associated methicillin resistance gene *mecA*, the PVL toxin and the toxic shock syndrome (TSS) toxin were not detected in any of the isolates **(Table 3)**. The immune evasion genes *scn*, *chp*, and *sak* were present in 23 (92%), 17 (68%) and 16 (64%) isolates, respectively **(Table 3)**. Further, the *sea* gene encoding for staphylococcal enterotoxin A, carried on an integrated prophage, was present in eight (32%) of the strains. There was no statistically significant difference in the presence/absence of these genes between mono- and co-infection strains. Unique prophage genes made up the greatest number of unique genes and were identified in 11 (44%) of strains, suggesting that phage integration is common.

**Table 3.**
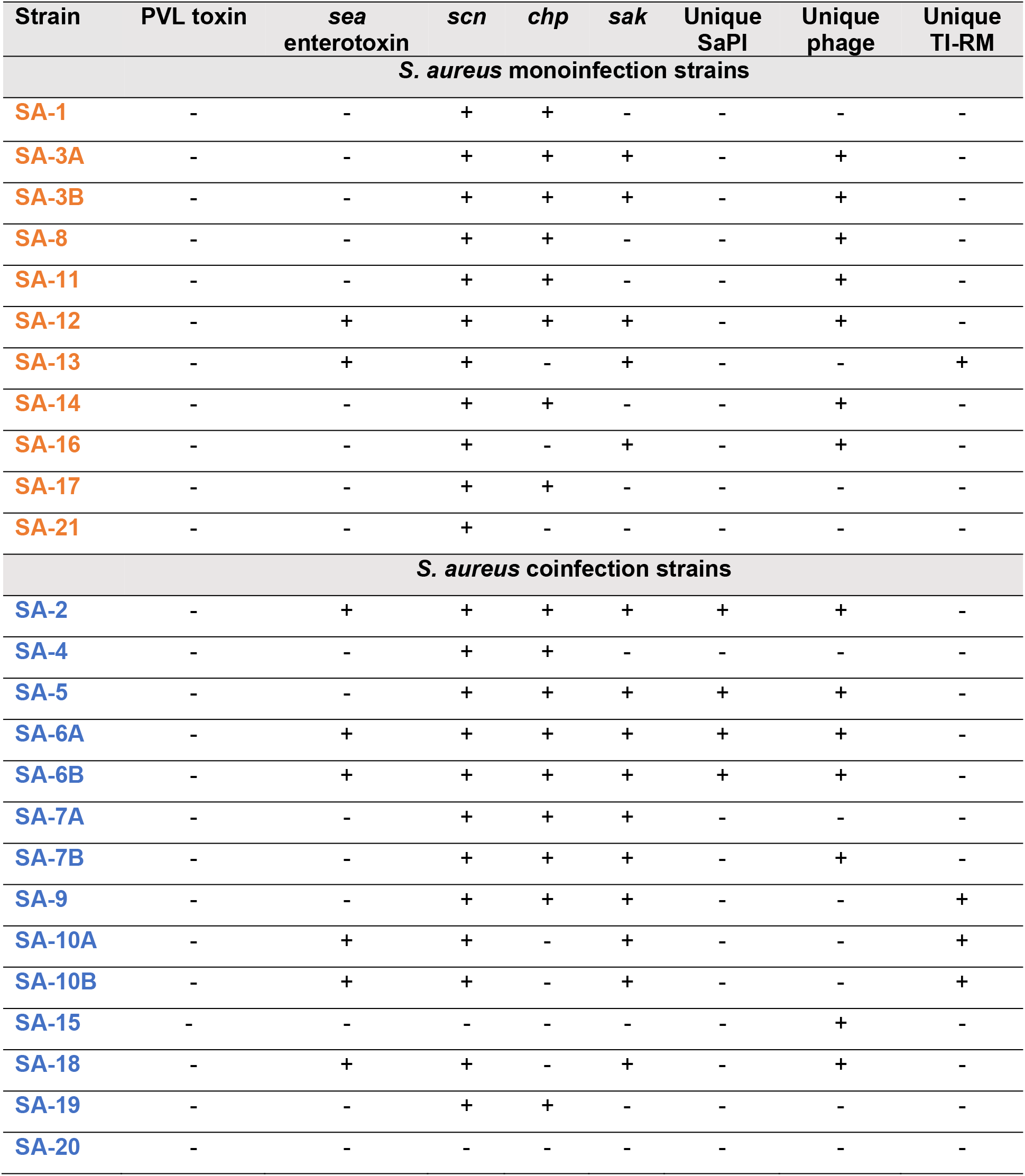
Presence or absence of unique genes relating to virulence in *S. aureus* clinical isolates. GenBank sequences of the clinical isolates and known reference genomes NCTC8325 and MRSA252 (NC_007795.1 and NC_002952.2, respectively) were aligned using progressive Mauve (21). Then, sequences were queried for the presence of genes likely to be related to virulence. Boxes populated with + indicate the gene is present and boxes populated with - indicate the gene is absent. The immune evasion genes *scn* encoding for the staphylococcal complement inhibitor*, chp* encoding for chemotaxis inhibitory protein, and *sak* encoding for staphylokinase were also examined. PVL, Panton Valentine leukocidin toxin; *sea*, staphylococcal enterotoxin A; SaPI, *S. aureus* pathogenicity islands; TI-RM, Type I restriction-modification enzymes.

Moreover, four of these strains (SA-2, SA-5, SA-6A, SA-6B) also contained unique SaPI genes, indicating that virulence factors are harboured on MGEs within the clinical strains. Of the 13 strains that did not contain unique phage genes, four (30%) contained unique copies of genes for Type I restriction-modification (TI-RM) enzymes (SA-13, SA-9, SA-10A, SA-10B). T-I RM systems are a known barrier to HGT as they prevent DNA uptake from different species and other *S. aureus* lineages, therefore it is possible they are restricting phage integration in those genomes (48).

The *agr* and *spa* type of each isolate was also characterised **(Figure 1)**. Four types of the *agr* accessory gene regulator system have been described (I-IV), and studies have reported *agr* type influence parthenogenesis and may impact clinical outcomes (49). The strains had a near equal split between *agr* type I (9, 36%) and type II (10, 40%), with a lower prevalence of type III strains (6, 25%) and none belonging to type IV, which is reportedly the rarest *agr* type (50, 51).

The varying numbers and base sequence of tandem repeats determines the *spa* type in *S. aureus* strains (52), which has been used to investigate *S. aureus* outbreaks. The isolates represented a wide range of *spa* types, and only isolates that were obtained from the same patient presented the same *spa* variant.

Average nucleotide identity calculations suggested that isolates collected from the same patients were likely the same strain (> 99.9). The maximum identity between isolates collected from two different patients was 99.98% (SA-13 and SA-15) and the minimum identity between any two clinical isolates was 97.34%. None of the isolates were closely related enough based on average nucleotide identity, *agr*, and *spa* type to suggest cross infection between patients.

### Haemolytic activity

Haemolysins are cytolytic toxins which are secreted by many *S. aureus* strains. They have been commonly recognised as essential virulence factors in murine CF infection models (53) and previous studies suggest strains downregulate expression of virulence factors during chronic infection (54). We therefore sought to establish whether our chronic infection isolates retained the ability to lyse ovine red blood cells (RBC), and whether there were any differences in haemolytic activity between the mono- and coinfection strains.

Eighteen (72%) of the isolates tested showed cytolytic activity against the RBC and there was no observable difference in lysis patterns between the monoinfection and coinfection *S. aureus* strains. However, interestingly we noted that four isolates that had lost their ability to lyse RBC came from patients that provided two separate *S. aureus* samples (3A, 6B, 7B and 10A), whilst the other patient sample retained its haemolytic activity **(Figure 3).** This is further evidence that multiple strains of the same species can exhibit different phenotypes within the same CF lung.

**Figure 3.**
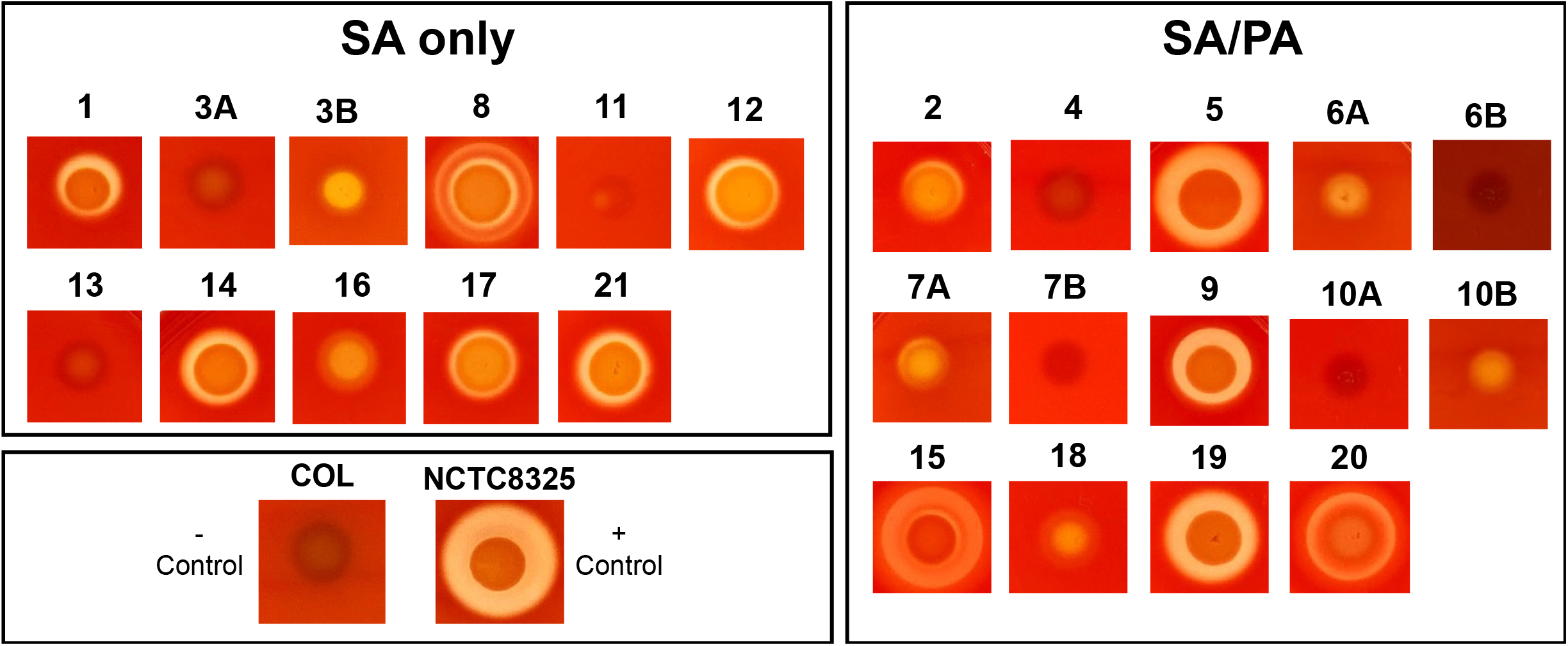
Haemolytic profiles of *S. aureus* isolates. *S. aureus* colonies were grown overnight in TSB (37 °C, 15 hrs), then 2 μl of each culture was spotted on to sheep blood agar plates. Cytolytic activity was recognised as clearance halos around the colonies. Colonies were spotted and plates incubated overnight at 37 °C followed by 4 °C incubation for 24 hours to detect hot-cold activation of b-haemolysin. COL and RN6390 were used as known negative and positive controls, respectively. Representative images from 3 biological replicates.

### Extracellular polysaccharide production

Another phenotype associated with chronic *S. aureus* infection is increased extracellular polysaccharide (EPS) production, which is an important constituent of biofilms (55). *S. aureus* produces one dominant EPS, chemically defined as poly-N-acetyl-β-(1–6)-glucosamine (PNAG) (55). PNAG is synthesised by proteins encoded at the intercellular adhesion (*icaABCD*) locus and its production is regulated by IcaR (56). PNAG overproduction could result in an advantageous mucoid phenotype as studies have found that mucoidy can increase intrinsic resistance to antibiotics, improve host immune-evasion and increase survival under nutrient limited conditions (15).

To establish whether any of the clinical isolates displayed a mucoid phenotype, EPS production was assayed by growth on Congo red agar plates. Most isolates had retained their ability to produce EPS (73%), which was also seen in a study of *S. aureus* isolates from chronically-infected CF individuals in the USA (49) **(Figure 4)**. Only 14% of isolates presented the overproducer mucoid colony morphology and 59% displayed normal EPS production. Separated by group, a higher number (4, 26%) of the coinfection strains presented the mucoid phenotype vs only 1 (9%) of the *S. aureus* monoinfection strains **(Figure 4)**, non-significant, using Fisher’s exact test (p=0.36).

**Figure 4.**
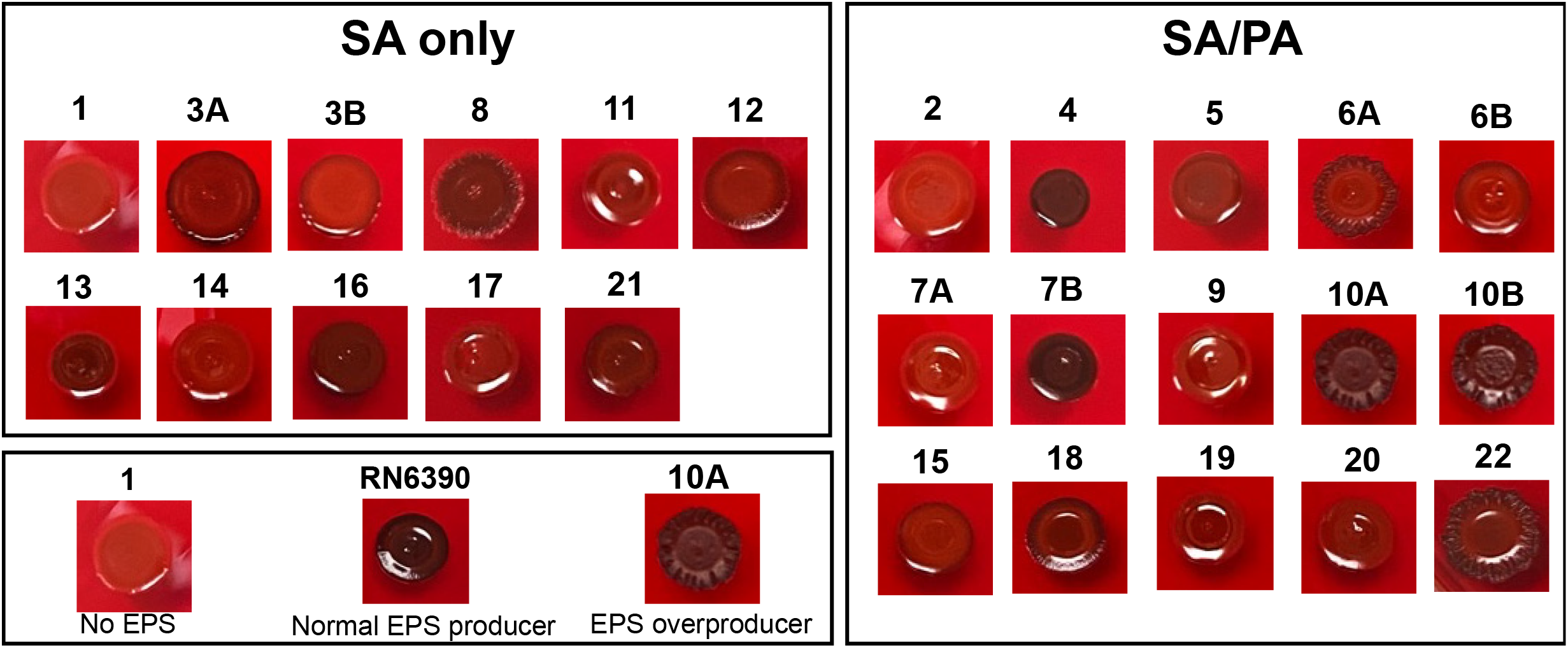
Majority of *S. aureus* clinical isolates retain their ability to produce EPS. *S. aureus* colonies were grown overnight in TSB (37 °C, 15 hrs) and 2 μl of each culture was spotted on to Congo red agar plates. Three phenotypes were revealed: EPS overproducers are dark and have a rough colony phenotype, normal EPS producers are dark in colour and appear shiny, and strains that lack all polysaccharide are bright red in colour and are also shiny. Representative images from 3 biological replicates.

Jefferson *et al*. identified a 5-nucleotide (TATTT) deletion in the *icaR-icaA* intergenic region that can induce over-expression of PNAG, and results in the mucoid phenotype (57). To establish whether the clinical isolates with a mucoid phenotype lacked this sequence, the DNA sequences in this region were queried. Two isolates which lacked the 5 bp nucleotide sequence exhibited the mucoid phenotype when tested (SA-10A, SA-10B). However, strains SA-6A and SA-8 did not present the deletion but exhibited a mucoid phenotype. Previous studies have suggested other unidentified mutations may also lead to this phenotype (50, 58). Additionally, although SA-6B and SA-7A lacked the 5 bp sequence, they presented a non-mucoid phenotype, this could be due to compensatory mutations elsewhere in the genome as no other mutations were detected in their *ica* loci. It is also noteworthy that the 5 bp nucleotide deletion was only observed in coinfection isolates, although this is non-significant using Fisher’s exact test (p=0.11).

### Antimicrobial resistance

It is being increasingly recognised that polymicrobial infections contribute to increased drug resistance of individual species (59). Previous work has identified modified drug resistance profiles in co-existing *S. aureus* and *P. aeruginosa* strains due to higher drug efflux pump activity and altered biofilm formation (60, 61). Therefore, to determine whether *S. aureus* strains isolated from *P. aeruginosa* coinfection presented increased antibiotic resistance, the resistance phenotypes of the *S. aureus* only and *S. aureus*/*P. aeruginosa* coinfection strains were assayed using the Kirby-Bauer disk diffusion method and the results are summarised in **Table 4**.

**Table 4.**
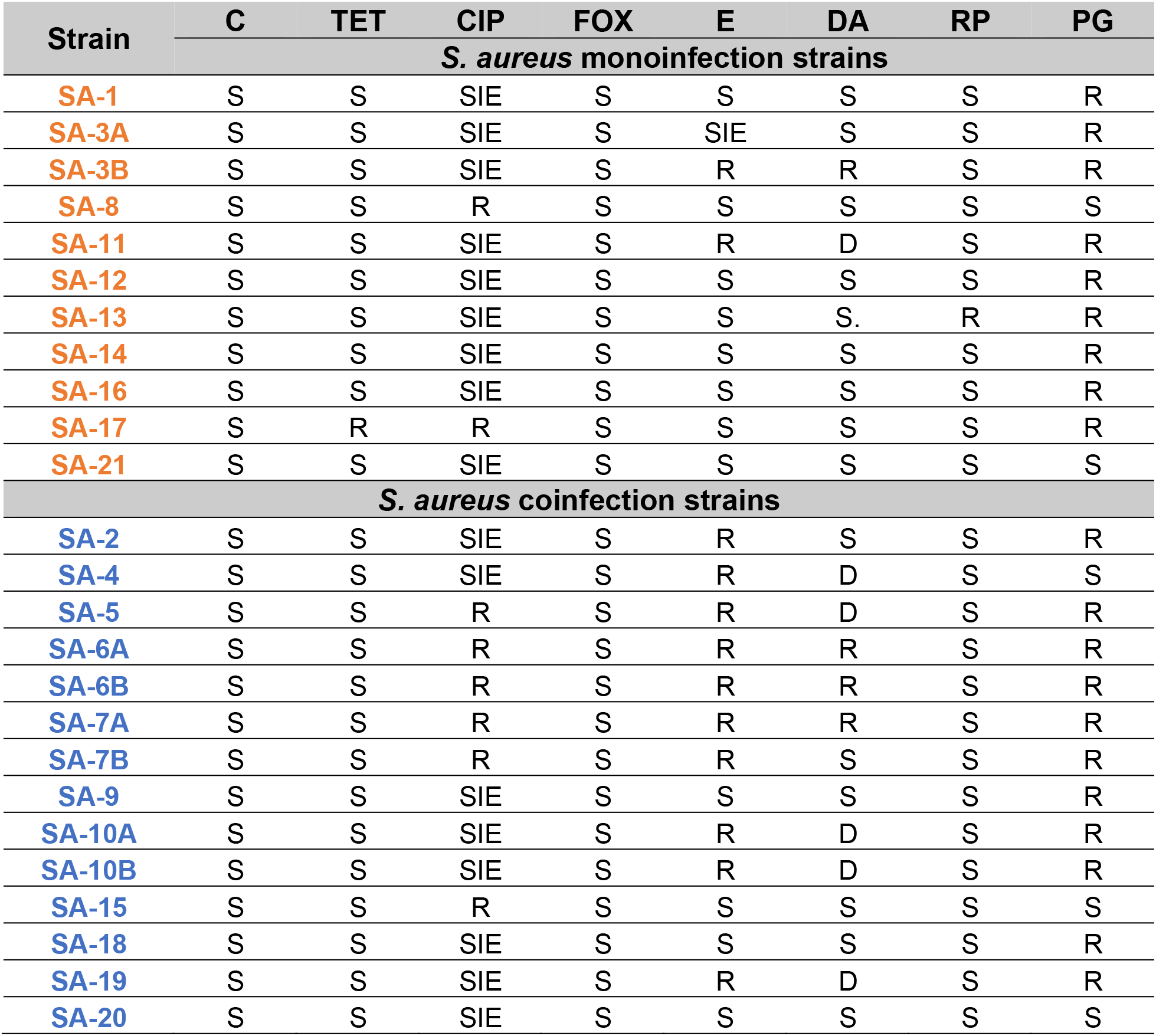
Antimicrobial drug resistance profiles of *S. aureus* clinical isolates. AST was assayed using the Kirby-Bauer disk diffusion method as outlined in the methods section. S-Susceptible. R-Resistant. SIE-Susceptible with increased exposure. D-Inducible clindamycin resistance. C-chloramphenicol. CIP-Ciprofloxacin. DA-Clindamycin. E-Erythromycin. FOX Cefoxitin. RP-Rifampicin. TET-Tetracycline. PG-Penicillin

| Strain | C | TET | CIP | FOX | E | DA | RP | PG |
| --- | --- | --- | --- | --- | --- | --- | --- | --- |
| ***S. aureus* monoinfection strains** |  |  |  |  |  |  |  |  |
| SA-1 | S | S | SIE | S | S | S | S | R |
| SA-3A | S | S | SIE | S | SIE | S | S | R |
| SA-3B | S | S | SIE | S | R | R | S | R |
| SA-8 | S | S | R | S | S | S | S | S |
| SA-11 | S | S | SIE | S | R | D | S | R |
| SA-12 | S | S | SIE | S | S | S | S | R |
| SA-13 | S | S | SIE | S | S | S. | R | R |
| SA-14 | S | S | SIE | S | S | S | S | R |
| SA-16 | S | S | SIE | S | S | S | S | R |
| SA-17 | S | R | R | S | S | S | S | R |
| SA-21 | S | S | SIE | S | S | S | S | S |
| ***S. aureus* coinfection strains** |  |  |  |  |  |  |  |  |
| SA-2 | S | S | SIE | S | R | S | S | R |
| SA-4 | S | S | SIE | S | R | D | S | S |
| SA-5 | S | S | R | S | R | D | S | R |
| SA-6A | S | S | R | S | R | R | S | R |
| SA-6B | S | S | R | S | R | R | S | R |
| SA-7A | S | S | R | S | R | R | S | R |
| SA-7B | S | S | R | S | R | S | S | R |
| SA-9 | S | S | SIE | S | S | S | S | R |
| SA-10A | S | S | SIE | S | R | D | S | R |
| SA-10B | S | S | SIE | S | R | D | S | R |
| SA-15 | S | S | R | S | S | S | S | S |
| SA-18 | S | S | SIE | S | S | S | S | R |
| SA-19 | S | S | SIE | S | R | D | S | R |
| SA-20 | S | S | SIE | S | S | S | S | S |

From the antimicrobial susceptibility phenotypes, there was some evidence that the coinfection strains showed increased resistance profiles compared to the *S. aureus* only strains **(Table 4)**, however this was non-significant (p=0.0562, X^2^=1.91,1). A significantly higher number of coinfection isolates (10, 71.4%) showed resistance to at least one type of macrolide, compared to only two monoinfection isolates (18.1%) using Fisher’s exact test, (p<0.0154). Additionally, a higher percentage of coinfection strains showed resistance to ciprofloxacin, an antibiotic commonly used with CF patients, compared to monoinfection strains (6, 42.8% vs 2, 18.1%), non-significant using Fisher’s exact test (p=0.2337). This is in line with previous studies showing that antibiotic pumps for ciprofloxacin are upregulated when *S. aureus* grows in coexistence with *P. aeruginosa* (61). Six coinfection isolates exhibited induced clindamycin resistance; this was established using the D-test, no monoinfection isolates presented induced clindamycin resistance, non-significant using Fisher’s exact test (*p* = 0.1804). Only one isolate was resistant to tetracycline (SA-17) and two isolates (SA-20 and SA-21) were sensitive to all antibiotics.

To identify the genes responsible for the resistance profile of the clinical isolates, the whole genome sequences were analysed using resistance prediction software ResFinder and MYKROBE **(Table S2)**. Comparing these predictions with our experimental testing, the programmes accurately predicted resistance profiles of 11 strains (52%). However, for the remaining isolates, they were predicted to be sensitive to macrolides (erythromycin and clindamycin), but antimicrobial susceptibility testing (AST) showed they exhibited resistance **(Table 3, Table S2)**. It is likely that the resistance gene may be carried on a plasmid rather than within the genome, as *ermC*, has been commonly identified on plasmids (62); plasmids were not selectively extracted in our study.

## Discussion

In this study we used a genotypic and phenotypic approach to characterise clinical *S. aureus* isolates and identify shared traits and differences between *S. aureus* isolates that coexist with *P. aeruginosa* vs *S. aureus* strains from patients who are *P. aeruginosa* negative. When comparing phylogenetic relationships and genotypes between strains, we noted that isolates of the same sequence type clustered together, however this did not correlate with the presence/absence of known virulence factors including haemolysin production, *sea* enterotoxin, or immune evasion genes. Additionally, in our study isolates did not group together based on their mucoid phenotype, as also noted by Bernardy *et al*. (50). However, isolates did largely cluster depending on whether they were isolated from mono- or coinfections. This data is also in accord with previous work (50), and hints at a possible genetic relationship between *S. aureus* isolates according to infection type.

Toxins and virulence factors are important for *S. aureus* pathogenicity, and the proportion of *S. aureus* strains encoding known virulence and toxin genes was high within our set of samples **(Table 3)**. Although previous reports have suggested that that *S. aureus* isolates adapted to the CF lung show reduced expression of α-haemolysin, most clinical isolates in this study retained their ability to lyse RBC, and there was no correlation between haemolytic phenotype and infection group.

The ability to produce EPS is an important factor in *S. aureus* persistence in the CF lung as it aids biofilm formation, provides protection from the host immune system, and increases tolerance to antibiotics (63, 64). Most isolates studied here were able to produce EPS when they were phenotypically characterised. However, the remainder were categorised as non-EPS producers and isolates that presented this phenotype belonged to both the mono- and coinfection groups. EPS is energetically costly to synthesise, and it is possible these isolates may instead take advantage of other EPS producers in the lung through formation of mixed-species biofilms. Although the EPS overproducer strains showed resistance to two or three different types of antibiotics, some of the non-producers were also multidrug resistant, therefore increased drug resistance profiles were not exclusively related to mucoidy.

Resistance to macrolides (erythromycin and clindamycin) was widespread among the strains with 12 (48%) strains showing resistance to this class of antibiotics, 9 of which belonged to the coinfection group (*p*<0.0154). This is interesting because although macrolides are not commonly used to combat *S. aureus* infections in the UK, long-term low-dose use of azithromycin is becoming a standard therapy for its anti-inflammatory properties and as a prophylactic in patients with chronic *P. aeruginosa* infection (2). Widespread azithromycin use in the UK may be driving increasing levels of *S. aureus* macrolide resistance, particularly in coinfection isolates as this treatment is more common in patients chronically infected with *P. aeruginosa*.

Previous studies have reported that 2-heptyl-4-hydroxyquinoline N-oxide, a quorum sensing signal produced by *P. aeruginosa*, provides *S. aureus* with protection against vancomycin (65). Additionally, transcriptome studies have shown *P. aeruginosa* induces expression of *S. aureus nor* genes, which encode drug efflux pumps for tetracycline and fluoroquinolone (61). However, no increased resistance against any of these antibiotics in coinfection isolates was observed, although it should be noted that Nor-dependent resistance may depend on close contact between the two organisms during testing (61) which was not carried out in our study.

All strains from patients that provided more than one isolate exhibited differing haemolytic abilities, and some also showed differing EPS production and antibiotic resistance phenotypes indicative of population heterogeneity within the same patient. This has previously been identified in CF sputum samples and it has been proposed that a more diverse community has increased viability and improved stress resistance (66).

In conclusion, *S. aureus* treatment and prophylaxis are greatly debated amongst clinicians, leading to differing treatment approaches across the world. A more detailed analysis of the nature of *S. aureus* CF isolates from different countries with different treatment regimes may ultimately allow for better informed and improved treatment of infections for people with CF.

## Acknowledgements

We are grateful to Newcastle University Molecular Microbiology MRes 4828F programme for supporting M.M. Authors acknowledge the support of Monash Bioinformatics Platform for this work.

## Ethical Statement

Respiratory samples were collected from patients attending the Adult Cystic Fibrosis Centre as part of their standard clinical care. Patients gave signed informed consent for the collection of clinical data for research and inclusion in the national CF registry, with approval from the regional ethics committee.

## Author contributions

MM and AP carried out all data collection and analysis, LR provided methodology for Pangenomic analysis. FRU carried out genome extractions, assembled the genomes and was involved in experimental planning and interpretation of results. SB and AP selected patients and collected appropriate samples. JHJ, LVB and NM sequenced the strains, assisted with whole genome assembly and bioinformatic analyses. TP and AYP conceptualised the study and TP and FRU supervised the study. MM, FRU, and TP drafted the manuscript. All authors edited the manuscript.

## References

1. Riordan JR, Rommens JM, Kerem B, Alon N, Rozmahel R, et al. Identification of the cystic fibrosis gene: cloning and characterization of complementary DNA. Science 1989;245(4922):1066–1073.

2. CF Trust UK. UK Cystic Fibrosis Registry Annual Data Report 2019.

3. Maliniak ML, Stecenko AA, McCarty NA. A longitudinal analysis of chronic MRSA and *Pseudomonas aeruginosa* co-infection in cystic fibrosis: A single-center study. J Cyst Fibros 2016;15(3):350–356.

4. Limoli DH, Yang J, Khansaheb MK, Helfman B, Peng L, et al. Staphylococcus aureus and *Pseudomonas aeruginosa* co-infection is associated with cystic fibrosis-related diabetes and poor clinical outcomes. Eur J Clin Microbiol Infect Dis 2016;35(6):947–953.

5. Rosenbluth DB, Wilson K, Ferkol T, Schuster DP. Lung function decline in cystic fibrosis patients and timing for lung transplantation referral. Chest 2004;126(2):412–419.

6. Briaud P, Bastien S, Camus L, Boyadjian M, Reix P, et al. Impact of Coexistence Phenotype Between *Staphylococcus aureus* and *Pseudomonas aeruginosa* Isolates on Clinical Outcomes Among Cystic Fibrosis Patients. Front Cell Infect Microbiol 2020;10:266.

7. Ahlgren HG, Benedetti A, Landry JS, Bernier J, Matouk E, et al. Clinical outcomes associated with *Staphylococcus aureus* and *Pseudomonas aeruginosa* airway infections in adult cystic fibrosis patients. BMC Pulm Med 2015;15:67.

8. Junge S, Gorlich D, den Reijer M, Wiedemann B, Tummler B, et al. Factors Associated with Worse Lung Function in Cystic Fibrosis Patients with Persistent Staphylococcus aureus. PLoS One 2016;11(11):e0166220.

9. Limoli DH, Hoffman LR. Help, hinder, hide and harm: what can we learn from the interactions between Pseudomonas aeruginosa and Staphylococcus aureus during respiratory infections? Thorax 2019;74(7):684–692.

10. Camus L, Briaud P, Vandenesch F, Doleans-Jordheim A, Moreau K. Mixed Populations and Co-Infection: *Pseudomonas aeruginosa* and Staphylococcus aureus. Adv Exp Med Biol 2022;1386:397–424.

11. Valentini M, Gonzalez D, Mavridou DA, Filloux A. Lifestyle transitions and adaptive pathogenesis of Pseudomonas aeruginosa. Curr Opin Microbiol 2018;41:15–20.

12. Goerke C, Wolz C. Adaptation of *Staphylococcus aureus* to the cystic fibrosis lung. Int J Med Microbiol 2010;300(8):520–525.

13. Tomlinson KL, Lung TWF, Dach F, Annavajhala MK, Gabryszewski SJ, et al. Staphylococcus aureus induces an itaconate-dominated immunometabolic response that drives biofilm formation. Nat Commun 2021;12(1):1399.

14. Hirschhausen N, Block D, Bianconi I, Bragonzi A, Birtel J, et al. Extended *Staphylococcus aureus* persistence in cystic fibrosis is associated with bacterial adaptation. Int J Med Microbiol 2013;303(8):685–692.

15. Schwartbeck B, Birtel J, Treffon J, Langhanki L, Mellmann A, et al. Dynamic in vivo mutations within the ica operon during persistence of Staphylococcus aureus in the airways of cystic fibrosis patients. PLoS Pathog 2016;12(11):e1006024.

16. Biswas L, Gotz F. Molecular Mechanisms of *Staphylococcus* and *Pseudomonas* Interactions in Cystic Fibrosis. Front Cell Infect Microbiol 2021;11:824042.

17. Graham BL, Steenbruggen I, Miller MR, Barjaktarevic IZ, Cooper BG, et al. Standardization of Spirometry 2019 Update. An Official American Thoracic Society and European Respiratory Society Technical Statement. Am J Respir Crit Care Med 2019;200(8):e70–e88.

18. Quanjer PH, Stanojevic S, Cole TJ, Baur X, Hall GL, et al. Multi-ethnic reference values for spirometry for the 3-95-yr age range: the global lung function 2012 equations. Eur Respir J 2012;40(6):1324–1343.

19. Wick RR, Judd LM, Gorrie CL, Holt KE. Unicycler: Resolving bacterial genome assemblies from short and long sequencing reads. PLoS Comput Biol 2017;13(6):e1005595.

20. Garrett SR, Mariano G, Palmer T. Genomic analysis of the progenitor strains of *Staphylococcus aureus* RN6390. Access Microbiol 2022;4(11): https://doi.org/10.1099/acmi.0.000464.v3

21. Darling AC, Mau B, Blattner FR, Perna NT. Mauve: multiple alignment of conserved genomic sequence with rearrangements. Genome Res 2004;14(7):1394–1403.

22. Seemann T. Prokka: rapid prokaryotic genome annotation. Bioinformatics 2014;30(14):2068–2069.

23. Edgar RC. MUSCLE: a multiple sequence alignment method with reduced time and space complexity. BMC Bioinformatics 2004;5:113.

24. Edgar RC. MUSCLE: multiple sequence alignment with high accuracy and high throughput. Nucleic Acids Res 2004;32(5):1792–1797.

25. Kumar S, Stecher G, Li M, Knyaz C, Tamura K. MEGA X: Molecular Evolutionary Genetics Analysis across Computing Platforms. Mol Biol Evol 2018;35(6):1547–1549.

26. Letunic I, Bork P. Interactive Tree Of Life (iTOL) v5: an online tool for phylogenetic tree display and annotation. Nucleic Acids Res 2021;49(W1):W293–W6.

27. Page AJ, Cummins CA, Hunt M, Wong VK, Reuter S, Holden MT, et al. Roary: rapid large-scale prokaryote pan genome analysis. Bioinformatics 2015;31(22):3691–3693.

28. Cantalapiedra CP, Hernandez-Plaza A, Letunic I, Bork P, Huerta-Cepas J. eggNOG-mapper v2: Functional Annotation, Orthology Assignments, and Domain Prediction at the Metagenomic Scale. Mol Biol Evol 2021;38(12):5825–5829.

29. Jolley KA, Bray JE, Maiden MCJ. Open-access bacterial population genomics: BIGSdb software, the PubMLST.org website and their applications. Wellcome Open Res 2018;3:124.

30. Francisco AP, Bugalho M, Ramirez M, Carrico JA. Global optimal eBURST analysis of multilocus typing data using a graphic matroid approach. BMC Bioinformatics 2009;10:152.

31. Raghuram V, Alexander AM, Loo HQ, Petit RA, 3rd, Goldberg JB, Read TD. Species-Wide Phylogenomics of the *Staphylococcus aureus agr* Operon Revealed Convergent Evolution of Frameshift Mutations. Microbiol Spectr 2022;10(1):e0133421.

32. Sanchez-Herrero JF. spaTyper: Staphylococcal protein A (*spa*) characterization pipeline. Zenodo. 2020.

33. Jain C, Rodriguez RL, Phillippy AM, Konstantinidis KT, Aluru S. High throughput ANI analysis of 90K prokaryotic genomes reveals clear species boundaries. Nat Commun 2018;9(1):5114.

34. Hunt M, Bradley P, Lapierre SG, Heys S, Thomsit M, et al. Antibiotic resistance prediction for *Mycobacterium tuberculosis* from genome sequence data with Mykrobe. Wellcome Open Res 2019;4:191.

35. Zankari E, Hasman H, Cosentino S, Vestergaard M, Rasmussen S, et al. Identification of acquired antimicrobial resistance genes. J Antimicrob Chemother 2012;67(11):2640–2644.

36. Carver T, Harris SR, Berriman M, Parkhill J, McQuillan JA. Artemis: an integrated platform for visualization and analysis of high-throughput sequence-based experimental data. Bioinformatics 2012;28(4):464–469.

37. Larkin MA, Blackshields G, Brown NP, Chenna R, McGettigan PA, et al. Clustal W and Clustal X version 2.0. Bioinformatics 2007;23(21):2947–2948.

38. Waterhouse AM, Procter JB, Martin DM, Clamp M, Barton GJ. Jalview Version 2--a multiple sequence alignment editor and analysis workbench. Bioinformatics 2009;25(9):1189–1191.

39. Burnside K, Lembo A, de Los Reyes M, Iliuk A, Binhtran NT, et al. Regulation of hemolysin expression and virulence of *Staphylococcus aureus* by a serine/threonine kinase and phosphatase. PLoS One 2010;5(6):e11071.

40. Feil EJ, Li BC, Aanensen DM, Hanage WP, Spratt BG. eBURST: inferring patterns of evolutionary descent among clusters of related bacterial genotypes from multilocus sequence typing data. J Bacteriol 2004;186(5):1518–1530.

41. Cao Z, Casabona MG, Kneuper H, Chalmers JD, Palmer T. The type VII secretion system of *Staphylococcus aureus* secretes a nuclease toxin that targets competitor bacteria. Nat Microbiol 2016;2:16183.

42. Ulhuq FR, Gomes MC, Duggan GM, Guo M, Mendonca C, et al. A membrane-depolarizing toxin substrate of the *Staphylococcus aureus* Type VII secretion system mediates intra-species competition. Proc Natl Acad Sci U S A 2020;17(34):20836–20847.

43. Bowman L, Palmer T. The Type VII Secretion System of Staphylococcus. Annu Rev Microbiol 2021;75:471–494.

44. Warne B, Harkins CP, Harris SR, Vatsiou A, Stanley-Wall N, et al. The Ess/Type VII secretion system of S*taphylococcus aureus* shows unexpected genetic diversity. BMC Genomics 2016;17(1):222.

45. Ibberson CB, Stacy A, Fleming D, Dees JL, Rumbaugh K, et al. Co-infecting microorganisms dramatically alter pathogen gene essentiality during polymicrobial infection. Nat Microbiol 2017;2:17079.

46. Haaber J, Penades JR, Ingmer H. Transfer of Antibiotic Resistance in Staphylococcus aureus. Trends Microbiol 2017;25(11):893–905.

47. McCarthy AJ, Witney AA, Lindsay JA. *Staphylococcus aureus* temperate bacteriophage: carriage and horizontal gene transfer is lineage associated. Front Cell Infect Microbiol 2012;2:6.

48. Oliveira PH, Touchon M, Rocha EP. The interplay of restriction-modification systems with mobile genetic elements and their prokaryotic hosts. Nucleic Acids Res 2014;42(16):10618–10631.

49. Abdulgader SM, van Rijswijk A, Whitelaw A, Newton-Foot M. The association between pathogen factors and clinical outcomes in patients with *Staphylococcus aureus* bacteraemia in a tertiary hospital, Cape Town. Int J Infect Dis 2020;91:111–118.

50. Bernardy EE, Petit RA, 3rd, Raghuram V, Alexander AM, Read TD, Goldberg JB. Genotypic and Phenotypic Diversity of *Staphylococcus aureus* Isolates from Cystic Fibrosis Patient Lung Infections and Their Interactions with *Pseudomonas aeruginosa*. mBio 2020;11(3).

51. Bibalan MH, Shakeri F, Javid N, Ghaemi A, Ghaemi EA. Accessory Gene Regulator Types of *Staphylococcus aureus* Isolated in Gorgan, North of Iran. J Clin Diagn Res 2014;8(4):DC07–DC09.

52. Mazi W, Sangal V, Sandstrom G, Saeed A, Yu J. Evaluation of *spa*-typing of methicillin-resistant *Staphylococcus aureus* using high-resolution melting analysis. Int J Infect Dis 2015;38:125–128.

53. Keitsch S, Riethmuller J, Soddemann M, Sehl C, Wilker B, et al. Pulmonary infection of cystic fibrosis mice with *Staphylococcus aureus* requires expression of alpha-toxin. Biol Chem 2018;399(10):1203–1213.

54. Camus L, Briaud P, Vandenesch F, Moreau K. How Bacterial Adaptation to Cystic Fibrosis Environment Shapes Interactions Between *Pseudomonas aeruginosa* and Staphylococcus aureus. Front Microbiol 2021;12:617784.

55. Nguyen HTT, Nguyen TH, Otto M. The staphylococcal exopolysaccharide PIA - Biosynthesis and role in biofilm formation, colonization, and infection. Comput Struct Biotechnol J 2020;18:3324–3334.

56. Heilmann C, Schweitzer O, Gerke C, Vanittanakom N, Mack D, Gotz F. Molecular basis of intercellular adhesion in the biofilm-forming *Staphylococcus epidermidis*. Mol Microbiol 1996;20(5):1083–1091.

57. Jefferson KK, Cramton SE, Gotz F, Pier GB. Identification of a 5-nucleotide sequence that controls expression of the *ica* locus in *Staphylococcus aureus* and characterization of the DNA-binding properties of IcaR. Mol Microbiol 2003;48(4):889–899.

58. Lennartz FE, Schwartbeck B, Dubbers A, Grosse-Onnebrink J, Kessler C, et al. The prevalence of *Staphylococcus aureus* with mucoid phenotype in the airways of patients with cystic fibrosis-A prospective study. Int J Med Microbiol 2019;309(5):283–287.

59. Nabb DL, Song S, Kluthe KE, Daubert TA, Luedtke BE, Nuxoll AS. Polymicrobial Interactions Induce Multidrug Tolerance in *Staphylococcus aureus* Through Energy Depletion. Front Microbiol 2019;10:2803.

60. Beaudoin T, Yau YCW, Stapleton PJ, Gong Y, Wang PW, et al. Staphylococcus aureus interaction with *Pseudomonas aeruginosa* biofilm enhances tobramycin resistance. NPJ Biofilms Microbiomes 2017;3:25.

61. Briaud P, Camus L, Bastien S, Doleans-Jordheim A, Vandenesch F, Moreau K. Coexistence with *Pseudomonas aeruginosa* alters *Staphylococcus aureus* transcriptome, antibiotic resistance and internalization into epithelial cells. Sci Rep 2019;9(1):16564.

62. Westh H, Hougaard DM, Vuust J, Rosdahl VT. Prevalence of *erm* gene classes in erythromycin-resistant *Staphylococcus aureus* strains isolated between 1959 and 1988. Antimicrob Agents Chemother 1995;39(2):369–373.

63. Donlan RM, Costerton JW. Biofilms: survival mechanisms of clinically relevant microorganisms. Clin Microbiol Rev 2002;15(2):167–193.

64. Vuong C, Voyich JM, Fischer ER, Braughton KR, Whitney AR, et al. Polysaccharide intercellular adhesin (PIA) protects *Staphylococcus epidermidis* against major components of the human innate immune system. Cell Microbiol 2004;6(3):269–275.

65. Orazi G, Jean-Pierre F, O’Toole GA. Pseudomonas aeruginosa PA14 Enhances the Efficacy of Norfloxacin against *Staphylococcus aureus* Newman Biofilms. J Bacteriol 2020;202(18):e00159–20.

66. Goerke C, Gressinger M, Endler K, Breitkopf C, Wardecki K, et al. High phenotypic diversity in infecting but not in colonizing *Staphylococcus aureus* populations. Environ Microbiol 2007;9(12):3134–3142.

