## Supplemental Tables for "Characterisation of key genotypic and phenotypic traits of clinical cystic fibrosis *Staphylococcus aureus* isolates"

| Mono-infection strains | Sample ID | Co-infection strains | Sample ID |
|---|---|---|---|
| SA-1 | ERS13476111 | SA-2 | ERS13476112 |
| SA-3A | ERS13476113 | SA-4 | ERS13476115 |
| SA-3B | ERS13476114 | SA-5 | ERS13476116 |
| SA-8 | ERS13476121 | SA-6A | ERS13476117 |
| SA-11 | ERS13476125 | SA-6B | ERS13476118 |
| SA-12 | ERS13476126 | SA-7A | ERS13476119 |
| SA-13 | ERS13476127 | SA-7B | ERS13476120 |
| SA-14 | ERS13476128 | SA-9 | ERS13476122 |
| SA-16 | ERS13476130 | SA-10A | ERS13476123 |
| SA-17 | ERS13476131 | SA-10B | ERS13476124 |
| SA-21 | ERS13476135 | SA-15 | ERS13476129 |
| | | SA-18 | ERS13476132 |
| | | SA-19 | ERS13476133 |
| | | SA-20 | ERS13476134 |

**Table S1. Clinical *S. aureus* strains under study with ENA sample ID accession numbers, project PRJEB56184.**

| Strain | Resistance genes | Predicted resistance phenotype | Observed resistance phenotype |
|---|---|---|---|
| | | ***S. aureus* monoinfection strains** | |
| SA-1 | *blaZ* | Penicillin resistant | Penicillin resistant |
| SA-3A | *blaZ* | Penicillin resistant | Penicillin resistant |
| SA-3B | *blaZ* | Penicillin resistant | Penicillin resistant<br>Erythromycin resistant<br>Clindamycin resistant |
| SA-8 | None | Sensitive to all antibiotics | Ciprofloxacin resistant |
| SA-11 | *blaZ*<br>*ermC* | Penicillin resistant<br>Erythromycin resistant<br>Clindamycin resistant | Penicillin resistant<br>Erythromycin resistant<br>D-Inducible Clindamycin resistance |
| SA-12 | *blaZ* | Penicillin resistant | Penicillin resistant |
| SA-13 | *blaZ*<br>*rpoB* H481N<br>*fusC* | Penicillin resistant<br>Rifampicin resistant<br>Fusidic acid resistant | Penicillin resistant<br>Rifampicin resistant<br>Fusidic acid not tested |
| SA-14 | *blaZ* | Penicillin resistant | Penicillin resistant |
| SA-16 | *blaZ* | Penicillin resistant | Penicillin resistant |
| SA-17 | *blaZ*<br>*tetK* | Penicillin resistant<br>Tetracycline resistant | Penicillin resistant<br>Tetracycline resistant<br>Ciprofloxacin resistant |
| SA-21 | None | Sensitive to all antibiotics | Sensitive to all antibiotics |
| | | ***S. aureus* coinfection strains** | |
| SA-2 | *blaZ* | Penicillin resistant | Penicillin resistant<br>Erythromycin resistant |
| SA-4 | *ermT* | Erythromycin resistant<br>Clindamycin resistant<br><br>Penicillin sensitive | Erythromycin resistant<br>Induced clindamycin resistance<br>Penicillin sensitive |
| SA-5 | *blaZ*<br>*grlA* E84K | Penicillin resistant<br>Ciprofloxacin resistant | Penicillin resistant<br>Ciprofloxacin resistant<br>Erythromycin resistant<br>D-Inducible Clindamycin resistance |
| SA-6A | *blaZ*<br>*grlA* E84K | Penicillin resistant<br>Ciprofloxacin resistant | Penicillin resistant<br>Ciprofloxacin resistant<br>Erythromycin resistant<br>Clindamycin resistant |
| SA-6B | *grlA* E84K | Penicillin sensitive<br>Ciprofloxacin resistant | Penicillin resistant<br>Ciprofloxacin resistant<br>Erythromycin resistant<br>Clindamycin resistant |
| SA-7A | *blaZ* | Penicillin resistant | Penicillin resistant<br>Ciprofloxacin resistant<br>Erythromycin resistant<br>Clindamycin resistant |

| | | | |
|---|---|---|---|
| **SA-7B** | *blaZ* | Penicillin resistant | Penicillin resistant<br>Ciprofloxacin resistant<br>Erythromycin resistant |
| **SA-9** | *blaZ* | Penicillin resistant | Penicillin resistant |
| **SA-10A** | *blaZ* | Penicillin resistant | Penicillin resistant<br>Erythromycin resistant<br>D-Inducible Clindamycin resistance |
| **SA-10B** | *blaZ* | Penicillin resistant | Penicillin resistant<br>Erythromycin resistant<br>D-Inducible Clindamycin resistance |
| **SA-15** | *fusC* | Fusidic acid resistant | Fusidic acid not tested<br>Ciprofloxacin resistant |
| **SA-18** | *blaZ*<br>*fusC* | Penicillin resistant<br>Fusidic acid resistant | Penicillin resistant<br>Fusidic acid not tested |
| **SA-19** | *blaZ*<br>*ermT* | Penicillin resistant<br>Erythromycin resistant<br>Clindamycin resistant | Penicillin resistant<br>Erythromycin resistant<br>D-Inducible Clindamycin resistance |
| **SA-20** | None | Sensitive to all antibiotics | Sensitive to all antibiotics |

**Table S2. Antibiotic resistance observed phenotypes vs predictions.** MYKROBE and ResFinder softwares were used to identify antimicrobial resistance genes within the genomes. FASTQ forward and reverse reads were submitted to MYKROBE (v.0.9.0) (34). FASTA sequences were uploaded to the online software ResFinder (v.4.1) (35). The observed phenotypes were then compared to the resistance phenotypes predicted by the softwares.
